# Autonomous AI-Driven Nanoscale Spatial Mapping Reveals Novel Targets and Ternary Architectures in 5xFAD Alzheimer’s Disease Model

**DOI:** 10.64898/2026.09.09.750448

**Authors:** M.V. Suissa, A. Koutures, S.A. Harker, B Vaughan, J Kfir, A Bhat, R Shihabi, M Dewal, E. Boyden, S. Taraman

## Abstract

Alzheimer’s disease (AD) is characterized by the deposition of amyloid-beta (Aβ) and microtubule-associated protein tau (MAPT) neurofibrillary tangles in the brain; however, the molecular mechanisms underlying associated synaptic dysfunction remain unclear. Eratos’ AI for Spatial Computing and Embedded Neurotherapeutic Discovery (ASCEND^TM^) engine was employed to analyze multiplexed expansion revealing (multiExR) data from the 5xFAD and wild-type mouse somatosensory cortex, enabling quantitative mapping of Aβ, RIM1, and GluA2 nanodomains. ASCEND^TM^ identified significant and previously unreported protein associations, including nanoscale colocalization of Aβ with postsynaptic AMPA-receptor subunit GluA2 and amyloid-bridged Aβ–GluA2–RIM1 ternary assemblies. These spatial signatures indicate complex synaptic disruption, defined by distinct morphological and density profiles associated with neurobiological and neuroinflammatory pathology. Integrating high-dimensional spatial computing with cross-modal literature synthesis advances traditional microscopy analysis toward autonomous, AI-driven scientific discovery. Identifying novel, druggable interfaces within native tissue expedites the discovery of precision neurotherapeutics for AD and other complex central nervous system disorders.

**Table of Contents Graphic**

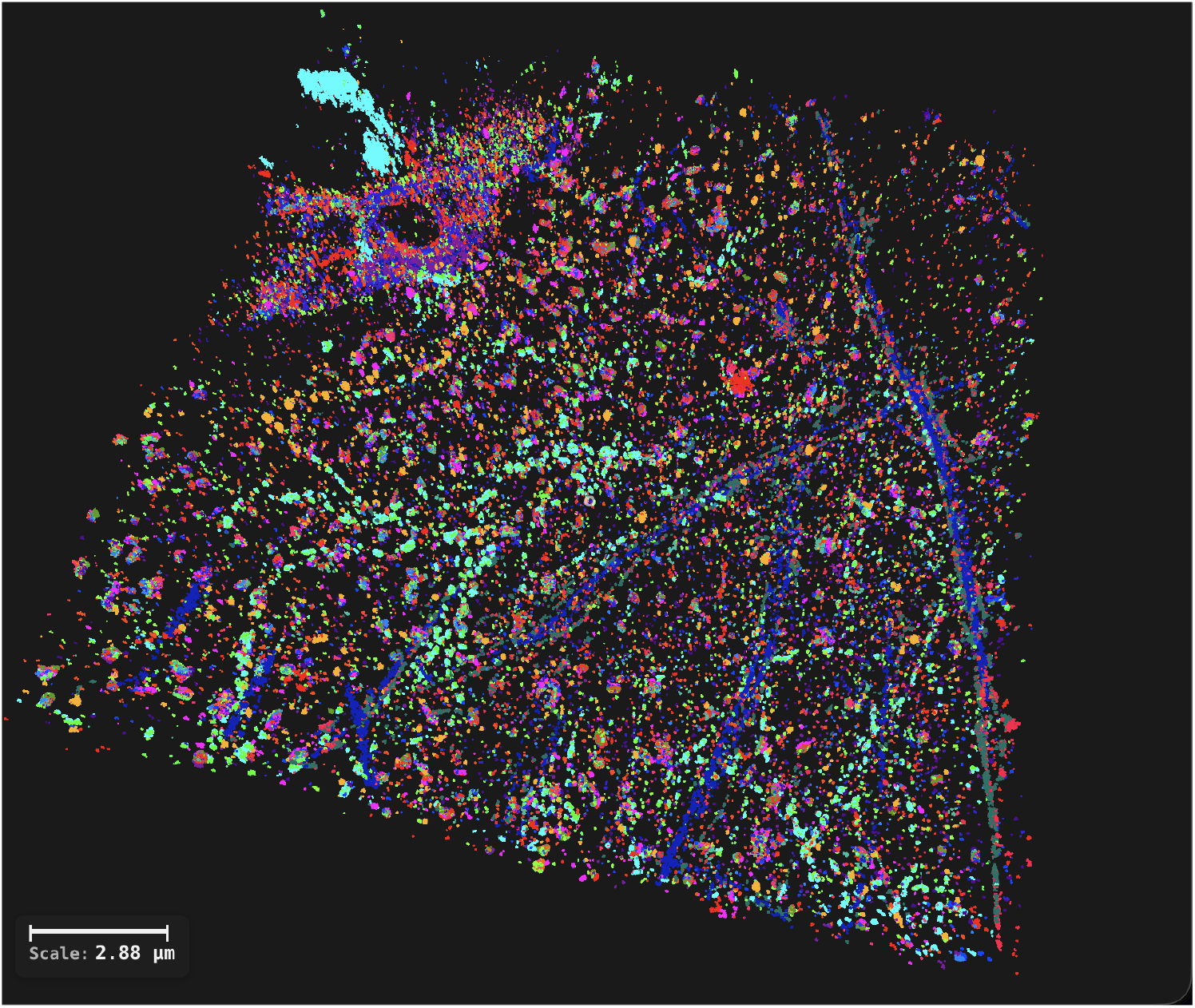

## Introduction

Alzheimer’s disease (AD) is clinically marked as progressive memory loss and cognitive impairment^1^. AD includes preclinical AD, prodromal AD, and dementia affecting approximately 416 million individuals worldwide^2^. Extracellular aggregations of amyloid beta (Aβ) plaques and resultant synaptic dysfunction are hallmarks of AD and are strongly associated with cognitive decline^3^. While protein abnormalities have been identified as a marker of the disease, elucidating the molecular underpinnings of synaptic dysfunction in AD remains a critical challenge^4^.

Though characterizing molecular associations with Aβ and AD-related synaptic dysfunction is difficult, several proteins have emerged as candidates. These include Rab3-interacting molecule 1 (RIM1) and α-amino-3-hydroxy-5-methyl-4-isoxazolepropionic acid receptor (AMPAR) subunit glutamate ionotropic receptor AMPA type subunit 2 (GluA2)^5–8^. RIM1 coordinates presynaptic active zone organization and synaptic vesicle docking, thus regulating neurotransmitter release activity^5^. AMPAR/ GluA2 governs postsynaptic excitatory synaptic dynamics^6^. Aβ oligomers have been implicated in weakening presynaptic function and inducing AMPAR trafficking deficits, particularly affecting GluA2-containing receptor complexes^7,8^. Despite recognition of these mechanistic links, direct nanoscale visualization of Aβ, RIM1, and GluA2 spatial relationships within brain tissue are limited by diffraction constraints of conventional microscopy.

The 5xFAD transgenic mouse model expresses five familial Alzheimer’s disease mutations and rapidly develops amyloid pathologies^9^. This mouse model phenotypically displays synaptic disruption that precedes overt neuronal death and involves impaired presynaptic vesicle release machinery and deregulated postsynaptic receptor function, making it ideal for studying AD-related synaptic abnormalities^8,10,11^. Expansion microscopy and its multiplexed variant (multiExR) improve nanoscale visualization by physically enlarging tissue specimens, enabling nanoscale imaging of multiple protein targets with nanoscale registration precision (∼25–40 nm) using conventional confocal microscopes^12^. This technology facilitates spatial reconstruction of molecular synaptic architectures and pathological aggregates simultaneously.

The ASCEND^TM^ engine was directed to analyze the raw multiExR data from Kang, J., Schroeder, M.E., Lee, Y. et al. (2024)^13^ to map novel, multidimensional spatial protein colocalizations. Double colocalizations refer to the spatial overlap or close proximity between two distinct protein nanodomains or molecular markers, reflecting direct or functionally coupled interactions^14^. This could include receptor clustering at the postsynaptic density or presynaptic active zone assembly^5,15^. Extending this concept, triple colocalizations denote the simultaneous spatial convergence of three distinct protein nanodomains, representing highly integrated molecular assemblies^16–17^. In this dataset, triple colocalizations involving Aβ, GluA2, and RIM1 indicate integrated synaptic microenvironments where amyloid converges with presynaptic and postsynaptic machinery. Such coordinated molecular alignment supports efficient synaptic transmission and facilitates rapid, activity-dependent synaptic plasticity fundamental to learning and memory^14,17^. Mapping these coupled nanodomains at high resolution provides critical insights into synaptic organization and dysfunction in health and disease^16–18^.

In this study, we demonstrate the capabilities of the ASCEND™ engine in surpassing traditional AI-assisted literature searches, facilitating a rapid closed-loop scientific discovery process that significantly accelerates the drug discovery pipeline. Our analysis identified 40,831 protein colocalizations across 9 5xFAD and 8 wild-type somatosensory mouse cortex regions, uncovering close spatial proximities between amyloid-beta (Aβ) and Rab3-interacting molecule 1 (RIM1), along with close colocalization between Aβ6E10 and glutamate ionotropic receptor AMPA type subunit 2 (GluA2) nanodomains. High-resolution mapping also highlighted new candidate interfaces, including triple colocalizations, suggesting coordinated disruptions among presynaptic and postsynaptic proteins in Alzheimer’s disease pathogenesis. By extending data interpretation from Kang, J., Schroeder, M.E., Lee, Y. et al.’s 2024 publication, we explored whether ASCEND™ could discover associations beyond human capability within existing expansion microscopy datasets of the 5xFAD mouse brain, focusing on the spatial organization of Aβ6E10 (via 6E10 antibody), RIM1, and GluA2, complemented with vascular and glial markers (Lectin+, SMI+, GFAP+). Analyzing raw multiExR data, the ASCEND™ engine revealed previously unreported molecular spatial features and associations, quantitatively characterizing nanocluster morphology and spatial proximity, and identifying thousands of biologically significant molecular clusters and nanospatial associations, all of which were further integrated with existing neurobiological literature.

## Results

### Nanocluster Morphology and Quantification

ASCEND^TM^ revealed extensive protein neighborhoods and nanoscale organization consistent with known synaptic structures, and segmented 161,678 distinct nanoclusters across Aβ, RIM1, GluA2, Lectin, and additional reference markers in the 5xFAD and WT scans (Fig. 1). Cluster volumes exhibited expected distributions, with Aβ6E10 clusters showing variable morphology relative to the more homogeneous RIM1 and GluA2 nanodomains (median volumes in 5xFAD: 291, 59, and 69 voxels respectively). Voxel size was calculated as 162.5 x 162.5 x 250 nm (post-expansion), corresponding to a pre-expansion voxel dimension of 9.03 × 9.03 × 13.89 nm, with the 18x expansion factor applied to all spatial dimensions (Fig.1). 5xFAD regions were compared to wild type to evaluate differences in likelihood of colocalizations.

**Figure 1:**
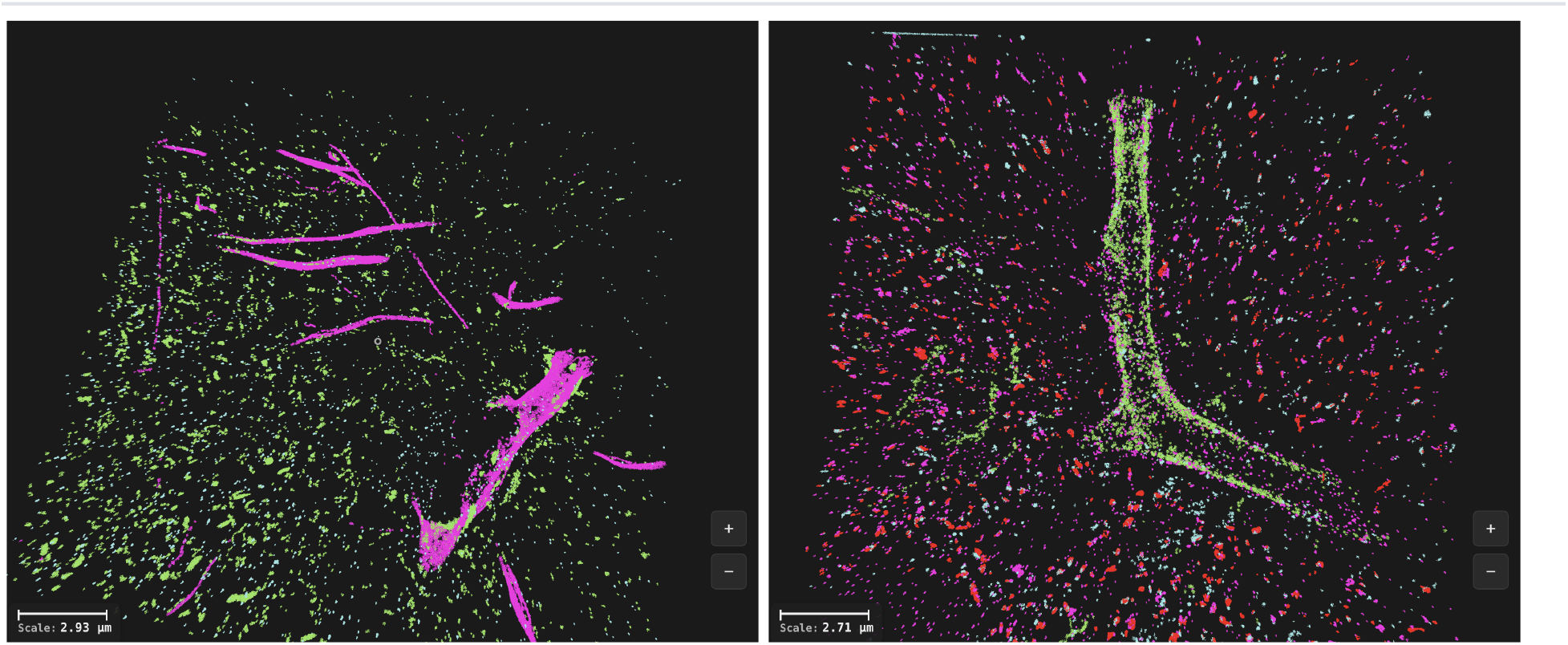
Wildtype and 5xFAD MultiExR Somatosensory Cortex Scans. Volumetric multiplexed expansion-revealing (multiExR) reconstructions from Kang et al. 2024 (Harvard Dataverse DOI:10.7910/DVN/JJBULY), processed with ASCEND. Left: WT mouse somatosensory cortex showing Lectin (magenta), RIM1 (green) and Aβ/Aβ6E10 (cyan). Right: 5xFAD mouse somatosensory cortex showing GluA2 (magenta), Lectin (green), Aβ6E10 (cyan) and RIM1 (red). Voxel size 162.5 × 162.5 × 250 nm post-expansion, corresponding to 9.03 × 9.03 × 13.89 nm pre-expansion at the 18× expansion factor reported by Kang et al. Clusters were segmented per channel by DBSCAN with voxel-aware ε on a p99.9 intensity threshold.

### Spatial Distribution and Co-localization of Nanoclusters

The distance metric refers to the average center-to-center distance between clusters, therefore, some colocalization distances are below the voxel resolution. More broadly, individual centroid distances below the multiExR registration uncertainty (∼25–40 nm) lie within the measurement limit and are reported as ‘within the registration/resolution limit’ rather than as exact molecular separations; only the distributions and the colocalization criterion (grid adjacency or ≤100 nm) are interpreted quantitatively. In the 5xFAD region, Aβ6E10 nanoclusters co-localized with RIM1 clusters at centroid-to-centroid distances as close as 8.05 nm (pre-expansion), with median Aβ–RIM1 separations of approximately 89.7 nm (pre-expansion). Notably, Aβ–RIM1 colocalization occurred at a rate of 1.4% in 5xFAD (Table 1; 2.67-fold enriched over a density-matched null, significant in 3 of 9 regions, Table 2), whereas no amyloid colocalization was detected in the WT condition (amyloid-negative control), highlighting a pathological association unique to the 5xFAD model (Table 1, 2). RIM1 and GluA2 clusters consistently formed spatially adjacent pairs, with a median centroid distance of 82.9 nm (pre-expansion, Table 3), consistent with their roles at presynaptic and postsynaptic sites forming functional synaptic nanodomains. GLUA1-RIM1 colocalization was significantly enriched in 5xFAD compared to WT (p = 0.008, Table 2). GluA2-RIM1 colocalization was significantly enriched in WT compared to 5xFAD by raw rate (15.1% vs 3.1%, Table 1), however the density-controlled enrichment did not differ significantly between genotypes (p = 0.093; Table 2), consistent with reduced synaptic abundance in the 5xFAD model. Spatial comparison of the RIM1-GluA2 pairwise colocalizations in the 5xFAD and WT lines demonstrated a preserved centroid distance, but significantly increased overlap in 5xFAD mice compared to WT (p=0.027, Table 3). Aβ and GluA2 were significantly co-localized in the 5xFAD mice (2.85-fold density-controlled enrichment, significant in 9 of 9 regions; Table 2). This was also the most frequent amyloid pairing by raw rate (93.9%; Table 1), although that high rate partly reflects the large size of amyloid clusters; the enrichment over a density-matched null provides the size-independent measure of association. Co-localization events were defined as nanocluster centroid pairs separated by ≤100 nm pre-expansion and/or clusters with overlapping voxel proximity, accounting for physical expansion. For each event, two metrics were computed: (1) grid overlap fraction, defined as the number of overlapping grid cells divided by the smaller cluster’s total cells; and (2) centroid-to-centroid distance between cluster averaged center points, converted to biological scale by dividing by the expansion factor. Within typical volumetric fields in the 5xFAD model, the observed colocalization rates were 1.4% for Aβ6E10–RIM1, 3.1% for GluA2–RIM1, and 93.9% for Aβ6E10–GluA2 (Table 1; per-pair enrichment and significance in Table 2). Extending the analysis to additional AMPA-receptor subunits, amyloid further associated with GluA1 and GluA3 as well as GluA2, with the strongest density-controlled enrichment for GluA1 (4.59-fold, significant in 8 of 9 regions; Table 2). The presynaptic GluA–RIM1 apposition was preserved across all subunits in 5xFAD (Table 2; GluA1 2.21-fold, GluA2 1.92-fold, GluA3 2.10-fold; each significant in at least 8 of 9 regions).

**Table 1:**
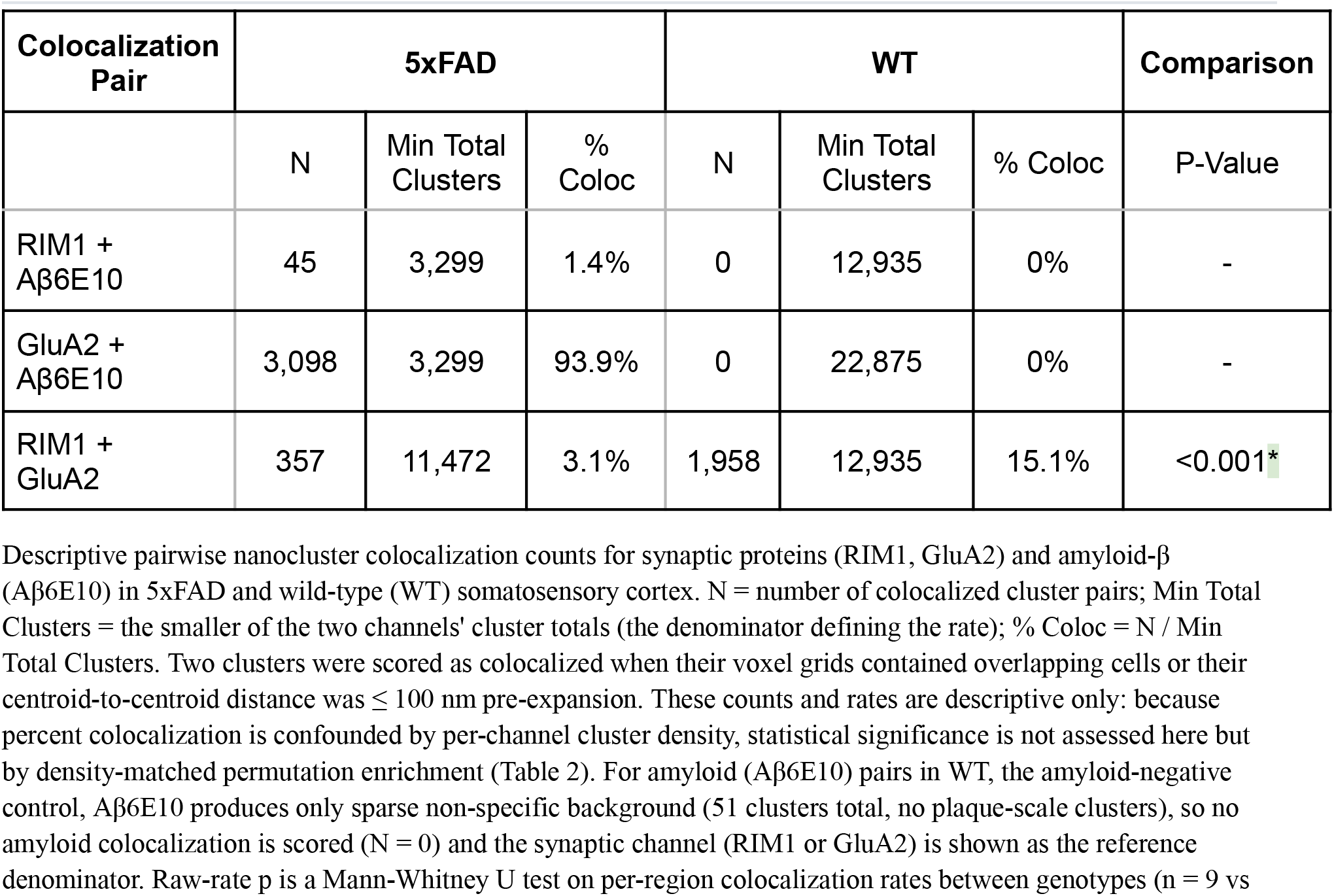
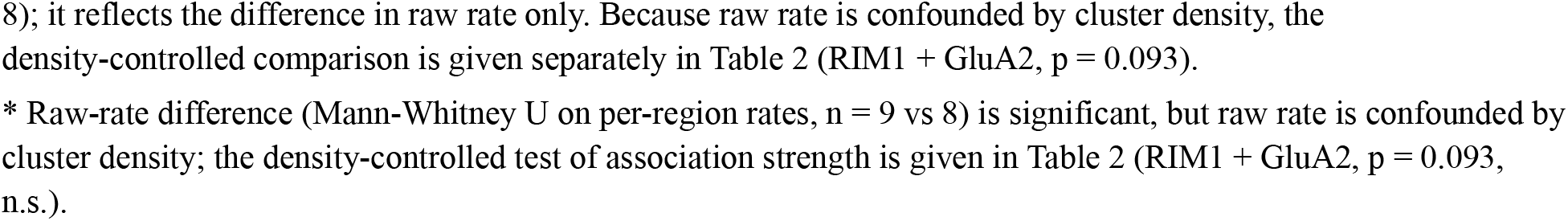
Pairwise Nanocluster Colocalization Rates In 5xFAD vs WT. Descriptive pairwise nanocluster colocalization counts for synaptic proteins (RIM1, GluA2) and amyloid-β (Aβ6E10) in 5xFAD and wild-type (WT) somatosensory cortex. N = number of colocalized cluster pairs; Min Total Clusters = the smaller of the two channels’ cluster totals (the denominator defining the rate); % Coloc = N / Min Total Clusters. Two clusters were scored as colocalized when their voxel grids contained overlapping cells or their centroid-to-centroid distance was ≤ 100 nm pre-expansion. These counts and rates are descriptive only: because percent colocalization is confounded by per-channel cluster density, statistical significance is not assessed here but by density-matched permutation enrichment (Table 2). For amyloid (Aβ6E10) pairs in WT, the amyloid-negative control, Aβ6E10 produces only sparse non-specific background (51 clusters total, no plaque-scale clusters), so no amyloid colocalization is scored (N = 0) and the synaptic channel (RIM1 or GluA2) is shown as the reference denominator. Raw-rate p is a Mann-Whitney U test on per-region colocalization rates between genotypes (n = 9 vs 8); it reflects the difference in raw rate only. Because raw rate is confounded by cluster density, the density-controlled comparison is given separately in Table 2 (RIM1 + GluA2, p = 0.093). * Raw-rate difference (Mann-Whitney U on per-region rates, n = 9 vs 8) is significant, but raw rate is confounded by cluster density; the density-controlled test of association strength is given in Table 2 (RIM1 + GluA2, p = 0.093, n.s.).

**Table 2:** Density-Controlled Enrichment & Significance. Density-matched permutation enrichment (observed colocalizations divided by a density-controlled label-shuffle null), computed per region with a per-pair deterministic seed so all values are exactly reproducible. Enrichment is the pooled ratio across regions; “Significant Regions” gives the number of regions in which the pair is significantly enriched (Benjamini-Hochberg FDR-corrected permutation, q < 0.05; significant regions reach the permutation-resolution floor of p < 0.002) out of 9 (5xFAD) or 8 (WT), so within-condition significance is read directly from this fraction. The between-genotype column is a Mann-Whitney U test on per-region enrichment ratios, computable only for pairs present in both conditions. Among these, GluA1 + RIM1 was significantly more enriched in 5xFAD than in WT (p = 0.008), whereas GluA2 + RIM1 (p = 0.093) and GluA3 + RIM1 (p = 0.963) did not differ between genotypes; for GluA2 + RIM1 the higher raw colocalization rate in WT (Table 1) is therefore attributable to cluster density rather than to a stronger association. A dash indicates not determined: amyloid (Aβ6E10) is the negative control in WT and is not assessed there.

| Colocalization Pair | 5xFAD |  | WT |  | Comparison |
| --- | --- | --- | --- | --- | --- |
|  | Enrichment | Significant Regions | Enrichment | Significant Regions | P-Value |
| RIM1 + Aβ6E10 | 2.67x | 3/9 | - | - | - |
| GLUA1 + Aβ6E10 | 4.59x | 8/9 | - | - | - |
| GluA2 + Aβ6E10 | 2.85x | 9/9 | - | - | - |
| GLUA3 + Aβ6E10 | 2.63x | 5/9 | - | - | - |
| RIM1 + GLUA1 | 2.21x | 9/9 | 2.03x | 8/8 | 0.008 |
| RIM1 + GluA2 | 1.92x | 8/9 | 2.11x | 8/8 | 0.093 |
| RIM1 + GLUA3 | 2.10x | 9/9 | 2.15x | 8/8 | 0.963 |

**Table 3:**
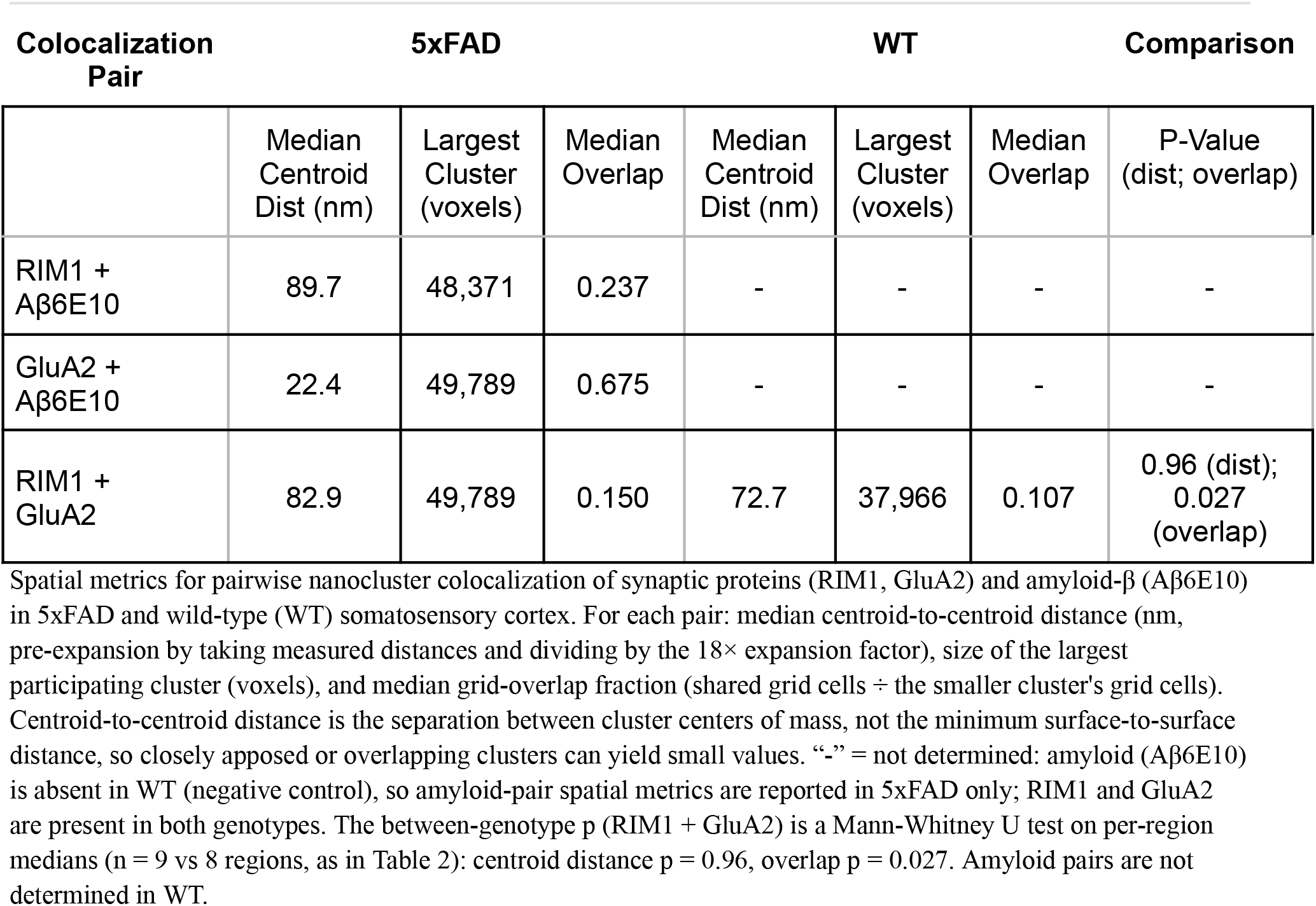
Spatial Properties of Pairwise Colocalizations. Spatial metrics for pairwise nanocluster colocalization of synaptic proteins (RIM1, GluA2) and amyloid-β (Aβ6E10) in 5xFAD and wild-type (WT) somatosensory cortex. For each pair: median centroid-to-centroid distance (nm, pre-expansion by taking measured distances and dividing by the 18× expansion factor), size of the largest participating cluster (voxels), and median grid-overlap fraction (shared grid cells ÷ the smaller cluster’s grid cells). Centroid-to-centroid distance is the separation between cluster centers of mass, not the minimum surface-to-surface distance, so closely apposed or overlapping clusters can yield small values. “-” = not determined: amyloid (Aβ6E10) is absent in WT (negative control), so amyloid-pair spatial metrics are reported in 5xFAD only; RIM1 and GluA2 are present in both genotypes. The between-genotype p (RIM1 + GluA2) is a Mann-Whitney U test on per-region medians (n = 9 vs 8 regions, as in Table 2): centroid distance p = 0.96, overlap p = 0.027. Amyloid pairs are not determined in WT.

After identifying colocalization events, we aimed to determine which specific nanocluster conformations drive these interactions. The foundational model of ASCEND^TM^ drives the spatial comparison process to select nanocluster subsets with a similarity threshold of ≥90% (Figure 2-3, Table 2). This model enables the isolation of the most structurally and molecularly homogeneous variants of RIM1, GluA2, and Aβ6E10 to those identified in colocalizations, emphasizing possible regions of highest colocalization (Fig. 3). By incorporating neighboring structural configurations, it performs colocalization analysis that not only controls for confounding heterogeneity but also identifies specific nanodomain conformations likely to interact spatially at biologically relevant scales. This spatial proximity is crucial for molecular interaction, allowing us to pinpoint critical molecular interfaces involved in synaptic pathology or physiology. This analysis allowed us to identify the number of RIM1 and GluA2 proteins that match the conformational state of those clustered with each other and Aβ6E10 (Fig. 3A-C). By distinguishing clusters of RIM1 and GluA2 with this precision, our model facilitates a detailed examination of presynaptic and postsynaptic nanoclusters, providing an accurate representation of canonical active zone and AMPA receptor assemblies.

**Figure 2:**
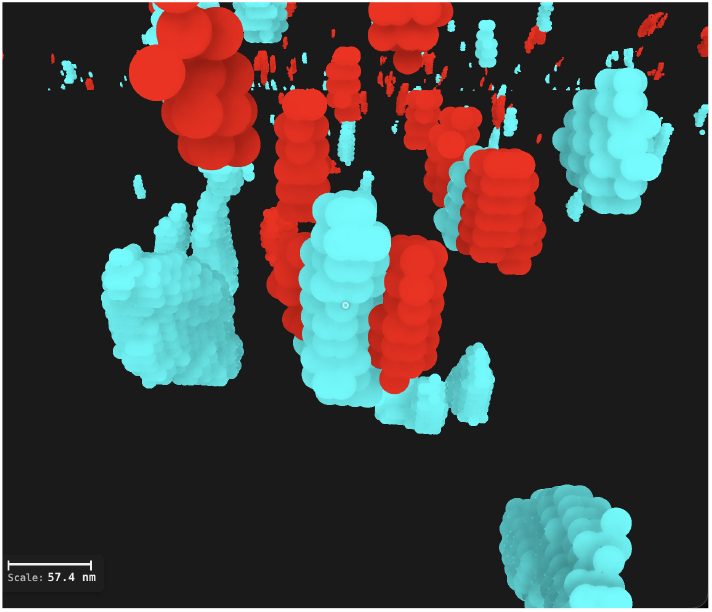
Representative 5xFAD Double-Colocalization Between RIM1 and Aβ6E10. ASCEND visualization of an individual RIM1–Aβ6E10 colocalization event in the 5xFAD somatosensory cortex (RIM1, red; Aβ6E10, cyan). Centroid-to-centroid distance 8.05 nm pre-expansion (within voxel resolution; see Fig. 1 caption); rendered at the pre-expansion voxel size given in Fig. 1.

**Figure 3:**
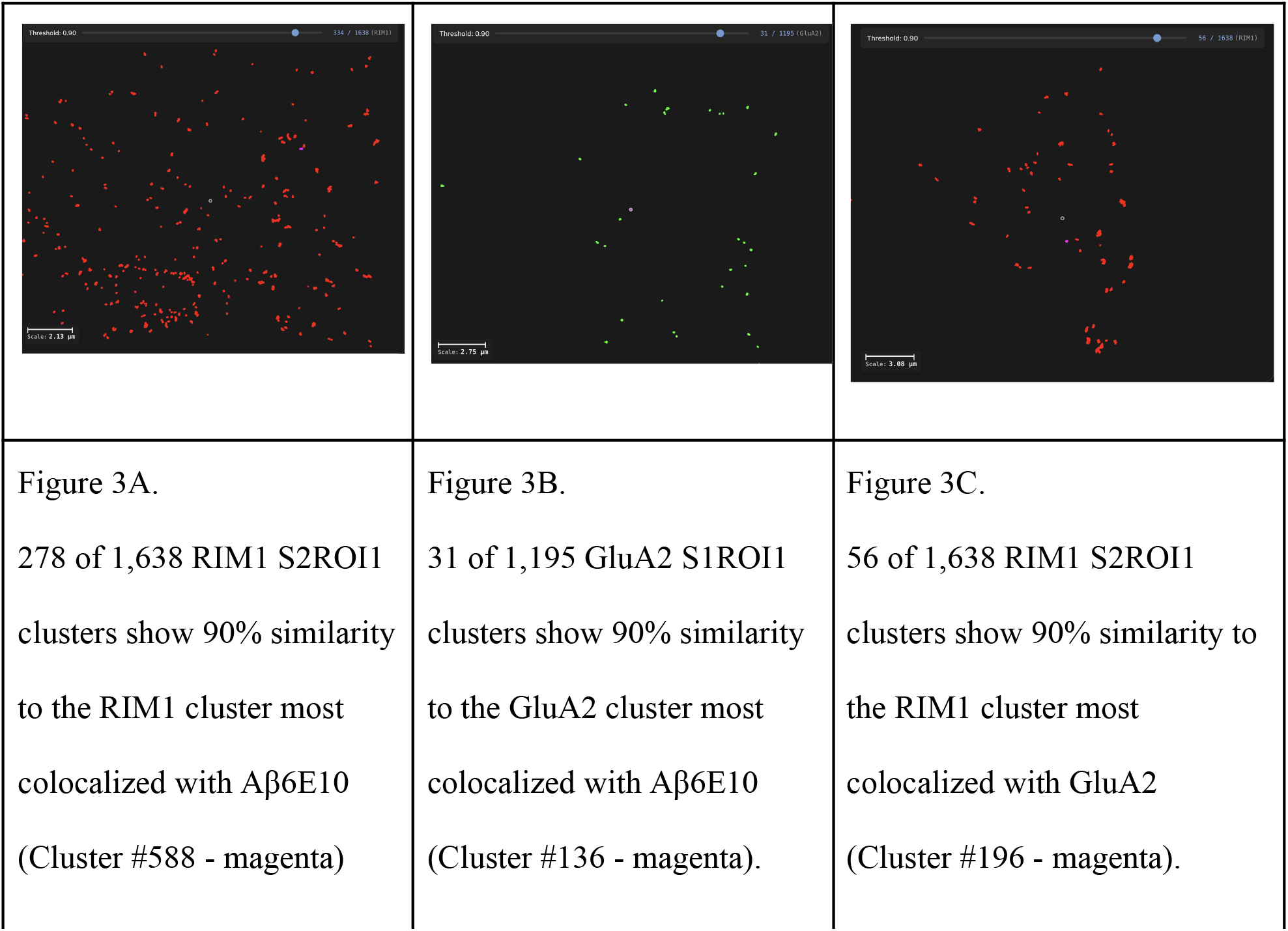
Structural Similarity Selection of Colocalized Cluster Sub-Populations. Our pre-trained PointJEPA embeddings were computed for every cluster in the indicated channel; cosine similarity was then evaluated against the cluster with the highest colocalization to the partner channel. Clusters at cosine ≥ 0.90 are highlighted. (A) 278 of 1,638 RIM1 clusters in S2ROI1 are ≥ 90% similar to RIM1 cluster #588 (magenta), the RIM1 cluster most strongly colocalized with Aβ. (B) 31 of 1,195 GluA2 clusters in S1ROI1 are ≥ 90% similar to GluA2 cluster #136 (magenta), the GluA2 cluster most strongly colocalized with Aβ. (C) 56 of 1,638 RIM1 clusters in S2ROI1 are ≥ 90% similar to RIM1 cluster #196 (magenta), the RIM1 cluster most strongly colocalized with GluA2.

### Triple Colocalizations

The triple colocalizations in the 5xFAD model involved GluA2 clusters with a median of 5,258 voxels, Aβ6E10 clusters with a median of 4,430 voxels, and RIM1 clusters with a median of 103 voxels, indicating that these ternary assemblies form preferentially among larger clusters. A total of 32 GluA2–RIM1–Aβ6E10 triple colocalizations were identified, localizing Aβ6E10 at key pre- and postsynaptic protein intersections (Tables 4 and 5, Figure 4). Conditional probability analysis quantifies how often a given pairwise colocalization sits inside the broader triple assembly: In 5xFAD, 71% of Aβ6E10–RIM1 events also recruited GluA2, compared with only 1% of the more abundant Aβ6E10–GluA2 events recruiting RIM1, indicating that amyloid completes the ternary assembly specifically at the rarer presynaptic contacts (Table 4). Because amyloid is absent in WT, this Aβ6E10–GluA2–RIM1 microdomain is unique to the 5xFAD model (Table 4). In total, 37 amyloid clusters each bridge a RIM1 and a GluA2 cluster, forming 53 ternary assemblies (Table 5). Of these 37 bridges, 32 are also closed triangles, meaning RIM1 and GluA2 are directly colocalized; the remaining 5 are bridge-only configurations in which Aβ6E10 is the sole link between the pre- and postsynaptic clusters. This bridge count exceeded the density-matched independence expectation (33.1 bridges) by 1.12-fold (permutation p = 0.013), indicating that amyloid bridging of RIM1 and GluA2 is modestly but significantly enriched beyond the product of its pairwise rates. The high conditional probability that an Aβ6E10–RIM1 contact also involves GluA2 (71%; Table 4) therefore largely reflects amyloid’s near-universal GluA2 association (∼90% of amyloid clusters), with a smaller specific ternary enrichment superimposed.

**Figure 4.**
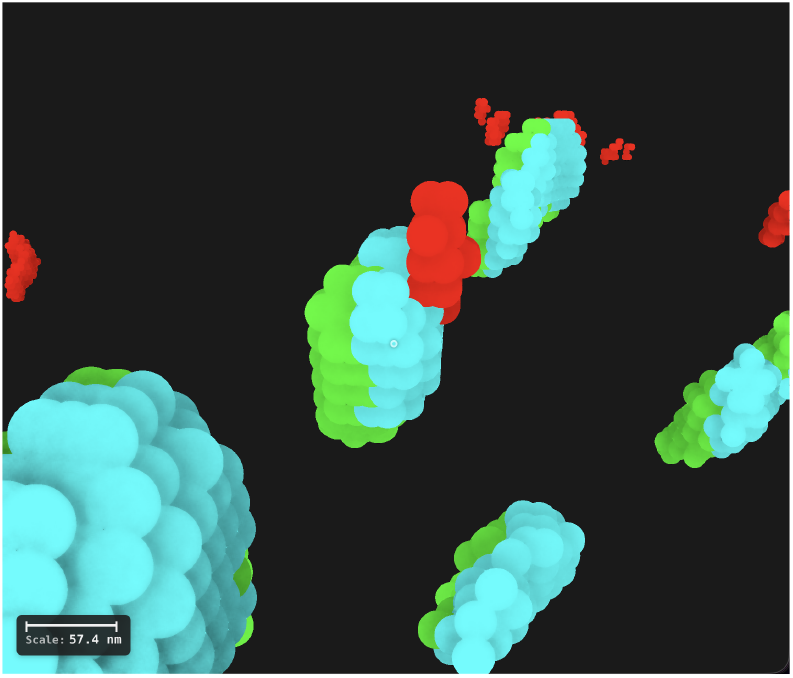
Representative 5xFAD Ternary Assembly (RIM1, GluA2, Aβ6E10) ASCEND visualization of a single GluA2 (magenta) – RIM1 (red) – Aβ6E10 (cyan) triple-colocalization event in 5xFAD somatosensory cortex.

**Table 4.**
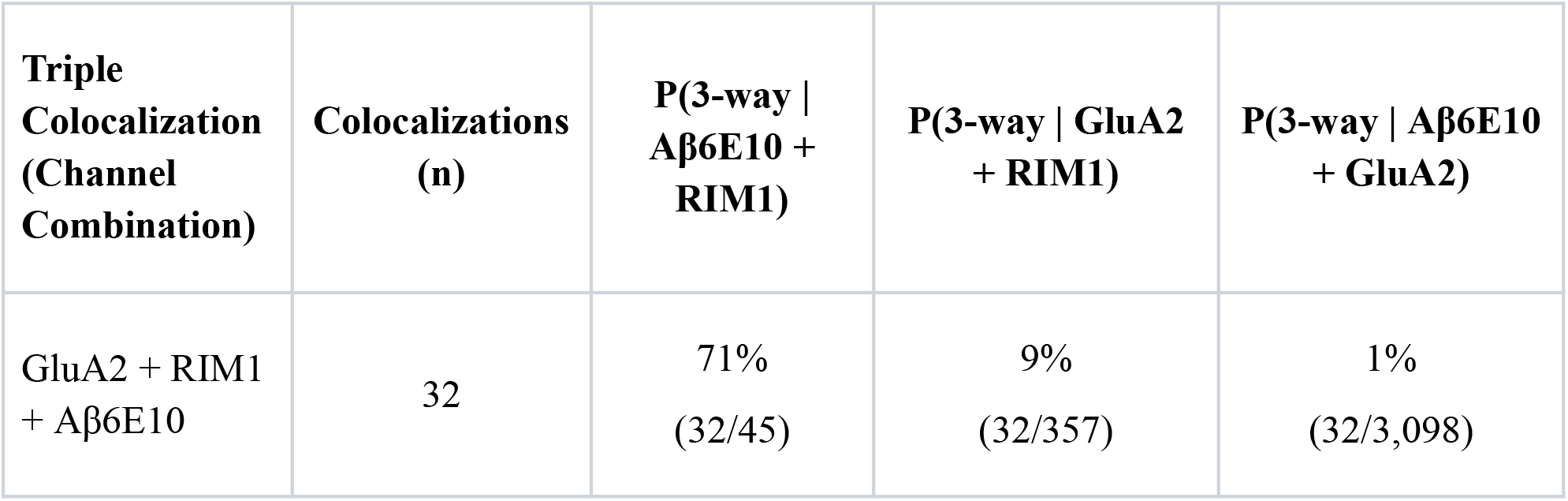
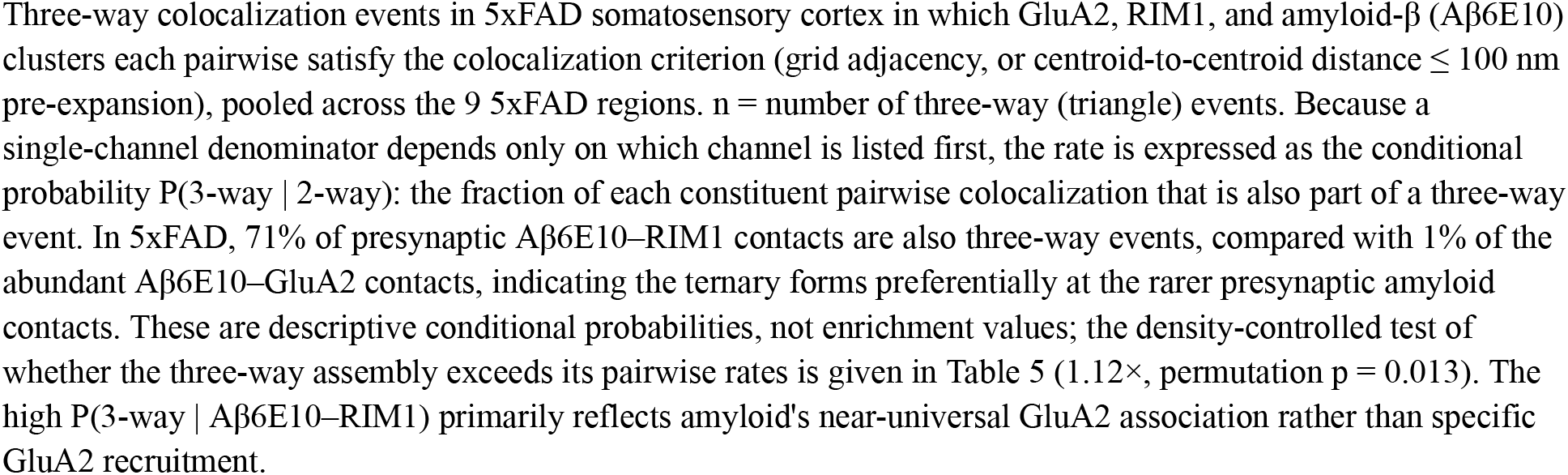
Three-Way Colocalization: GluA2–RIM1–Aβ6E10 - 5xFAD. Three-way colocalization events in 5xFAD somatosensory cortex in which GluA2, RIM1, and amyloid-β (Aβ6E10) clusters each pairwise satisfy the colocalization criterion (grid adjacency, or centroid-to-centroid distance ≤ 100 nm pre-expansion), pooled across the 9 5xFAD regions. n = number of three-way (triangle) events. Because a single-channel denominator depends only on which channel is listed first, the rate is expressed as the conditional probability P(3-way | 2-way): the fraction of each constituent pairwise colocalization that is also part of a three-way event. In 5xFAD, 71% of presynaptic Aβ6E10–RIM1 contacts are also three-way events, compared with 1% of the abundant Aβ6E10–GluA2 contacts, indicating the ternary forms preferentially at the rarer presynaptic amyloid contacts. These are descriptive conditional probabilities, not enrichment values; the density-controlled test of whether the three-way assembly exceeds its pairwise rates is given in Table 5 (1.12×, permutation p = 0.013). The high P(3-way | Aβ6E10–RIM1) primarily reflects amyloid’s near-universal GluA2 association rather than specific GluA2 recruitment.

**Table 5.**
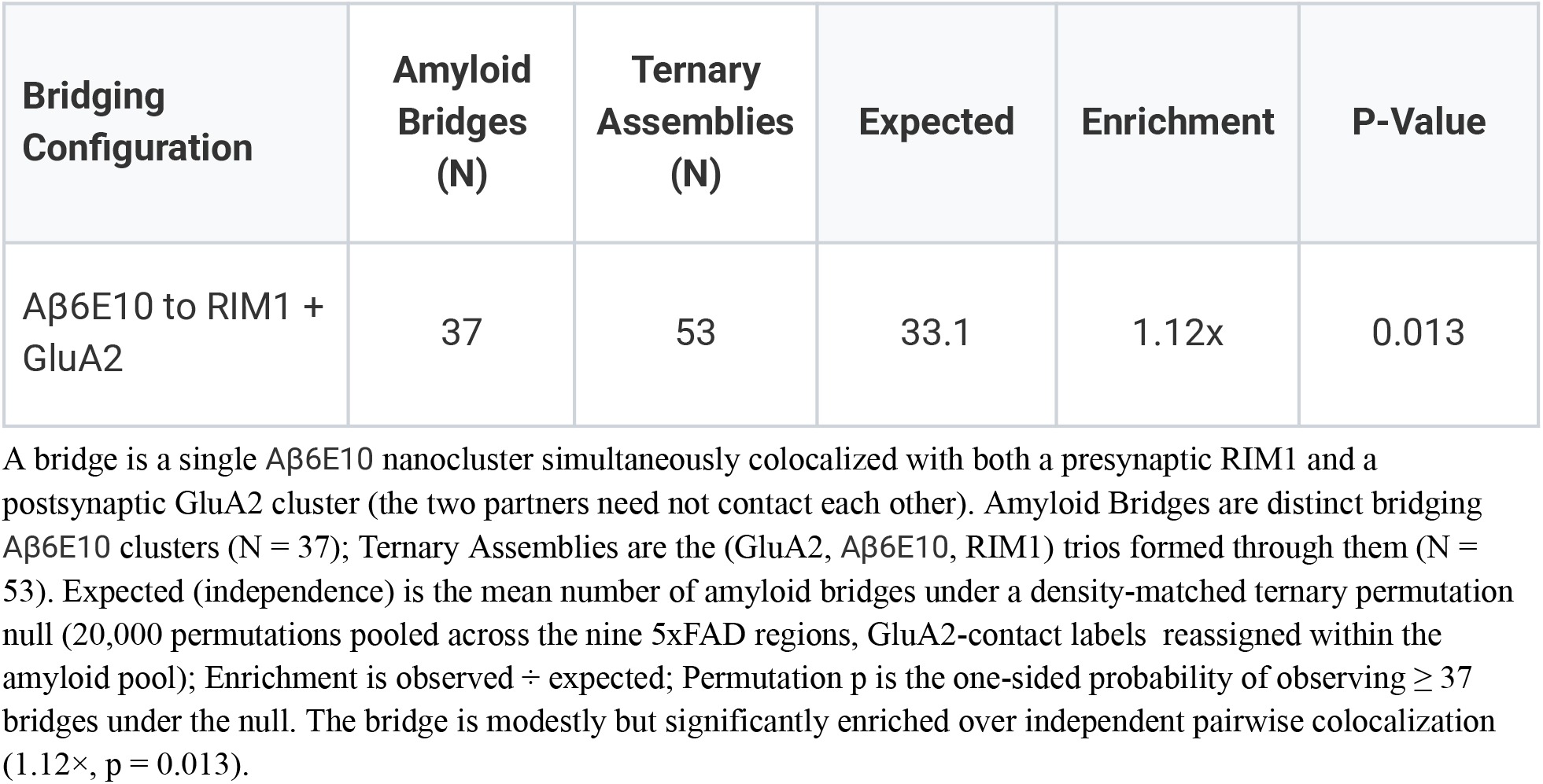
Three-Way N-Dimensional Colocalizations (NDCs): Amyloid Bridging RIM1 + GluA2 (5xFAD) A bridge is a single Aβ6E10 nanocluster simultaneously colocalized with both a presynaptic RIM1 and a postsynaptic GluA2 cluster (the two partners need not contact each other). Amyloid Bridges are distinct bridging Aβ6E10 clusters (N = 37); Ternary Assemblies are the (GluA2, Aβ6E10, RIM1) trios formed through them (N = 53). Expected (independence) is the mean number of amyloid bridges under a density-matched ternary permutation null (20,000 permutations pooled across the nine 5xFAD regions, GluA2-contact labels reassigned within the amyloid pool); Enrichment is observed ÷ expected; Permutation p is the one-sided probability of observing ≥ 37 bridges under the null. The bridge is modestly but significantly enriched over independent pairwise colocalization (1.12×, p = 0.013).

### Morphometric Differences Correlated with Spatial Relationships

To distinguish genuine changes in molecular association from differences in cluster abundance, we compared colocalization across all 9 5xFAD and 8 WT regions using density-matched permutation enrichment, which controls for per-channel cluster density (Table 2). This analysis showed that several apparent genotype differences in raw colocalization rate were largely attributable to differences in cluster abundance rather than to changes in association strength. GluA2–RIM1 colocalization occurred at a higher raw rate in WT than in 5xFAD (15.1% vs 3.1%; Table 1), yet the density-controlled enrichment did not differ significantly between genotypes (2.11-fold vs 1.92-fold; p = 0.093; Table 2), consistent with a reduction in GluA2 cluster abundance in 5xFAD (12,339 vs 22,875 clusters), rather than a loss of pre- and postsynaptic apposition. The synaptic GLUA–RIM1 association was preserved across AMPA-receptor subunits in 5xFAD and was in fact significantly stronger for GLUA1 (2.21-fold vs 2.03-fold; p = 0.008; Table 2). In contrast, Aβ6E10 associations were unique to the 5xFAD model, as amyloid was effectively absent in WT (51 background clusters, no plaque-scale clusters). Together, these results indicate that the principal disease-associated change is the emergence of amyloid–synaptic associations superimposed on a largely preserved organization of presynaptic and postsynaptic AMPA-receptor nanodomains, with apparent reductions in synaptic colocalization rate reflecting decreased synaptic abundance rather than disrupted molecular apposition (Tables 1 and 2).

### Discovery of Novel Protein Spatial Associations Through Literature Integration

Analysis of raw multiExR data uncovered thousands of previously unreported protein co-localization and spatial adjacency events beyond conventional interpretive scope. Notably, multiscale motifs involving Aβ6E10, RIM1, GluA2, and additional synaptic constituents were revealed, expanding the known molecular landscape of synaptic nanostructure alterations in the 5xFAD and WT somatosensory cortex (Figures 1-2, Tables 1-2). Filtering these colocalizations against the comprehensive neurobiological literature across scientific databases, ASCEND™ matched 1,678 to documented protein associations, 727 of which involved amyloid-β with synaptic proteins and were aligned with well-established Alzheimer’s disease pathology, while the remaining 951 represented novel spatial links not previously reported, implicating unexplored molecular participants and potential regulatory nodes within synaptic nanodomains.

### Autonomous Experimental Design and Drug Development Strategy Through Mechanistic Modeling and Target Identification

ASCEND^TM^ subsequently formulated a comprehensive experimental roadmap incorporating high-resolution spatial imaging to validate targeted nanocluster dynamics, optogenetic functional assays to assess synaptic efficacy, and the design of ligand libraries optimized for binding within spatially constrained active sites. Its integrated mechanistic models predicted several novel druggable interfaces that spatially coincide with amyloid nanoclusters and synaptic active zone proteins. Among these, specific allosteric modulation sites on RIM1-related complexes and GluA2-associated pathways emerged as prime candidates for therapeutic intervention to restore synaptic transmission fidelity (Figure 5).

**Figure 5.**
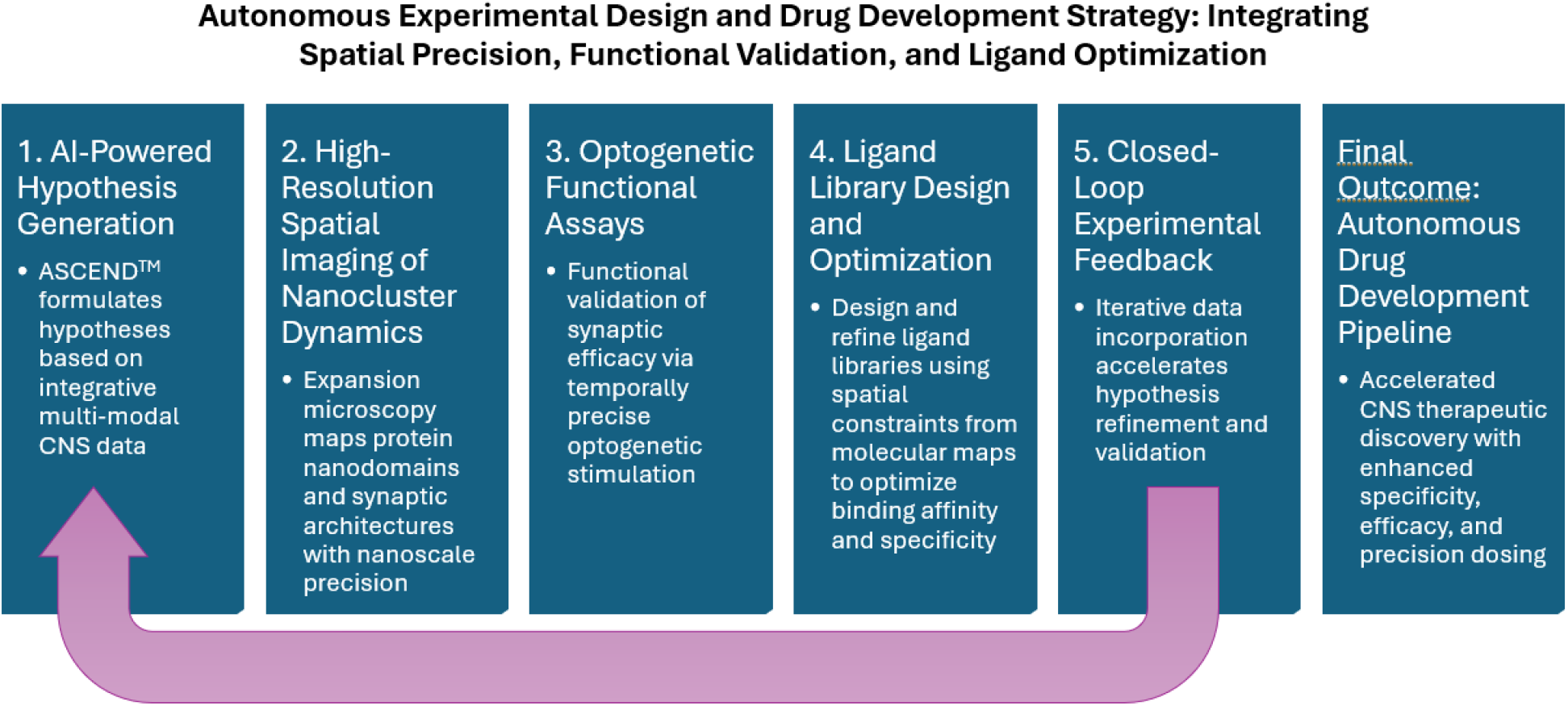
Comprehensive Experimental Roadmap. Pathway to demonstrate comprehensive experimental roadmap incorporating high-resolution spatial imaging to validate targeted nanocluster dynamics.

## Discussion and Conclusions

The current study utilized ASCEND^TM^ nanoscale spatial mapping with multiExR data to discover critical insights into the disease emergent synaptic pathology of the 5xFAD mouse model of AD. Advanced spatial processing allowed us to observe a previously unreported ternary nDC formed by Aβ6E10-GluA2–RIM1, and elucidate the mechanism by which Aβ6E10 selectively exploits a cryptic pocket at the transsynaptic interface. The ASCEND^TM^ analysis resulted in our ability to identify a druggable interface for potential mediation of synaptic dysfunction in AD.

We observed that GluA1–RIM1 association was significantly enriched in the 5xFAD model relative to wild-type (2.21× vs 2.03×; p = 0.008; Table 2), consistent with previously reported increases in GluA1-containing AMPAR signaling in AD^19^. Importantly, density control distinguished genuine association changes from differences in cluster abundance. GluA2–RIM1 showed a higher raw colocalization rate in WT than in 5xFAD samples (15.1% vs 3.1%; Table 1), however density-controlled enrichment revealed no significant difference between genotypes (1.92× vs 2.11×; p = 0.093; Table 2). This indicates that the lower GluA2–RIM1 rate in 5xFAD reflects reduced GluA2 cluster abundance (12,339 vs 22,875 clusters), a consequence of synaptic loss rather than a disruption of presynaptic to postsynaptic apposition.

ASCEND^TM^ analysis further detected subtle nanocluster morphological differences beyond spatial proximity, revealing features that are not easily discernible through human inspection, including size differences (GluA2 cluster size ∼69 voxels, Aβ6E10 cluster size ∼291 voxels), shape metrics, and neighborhood statistical context. We found that Aβ6E10 nanoclusters lie in close spatial proximity to RIM1 nanodomains with an average distance of 89.70 nm (pre-expansion), suggesting direct or indirect disruption of presynaptic machinery (<100nm)^19^. Previous reports indicate impaired vesicle docking and neurotransmitter release in amyloid-rich environments^20^, however the spatial heterogeneity of RIM1-Aβ6E10 nanoclusters implies localized synaptic vulnerability, potentially reflecting stages of amyloid plaque formation or regional synaptic resilience.

GluA2-containing AMPAR maintained close adjacency to RIM1 and was strongly colocalized with Aβ6E10 aggregates (93.9%; the most frequent amyloid pairing, Tables 1–3). These findings support a model where postsynaptic receptor architecture is directly engaged by amyloid, alongside indirect contributions from soluble oligomeric Aβ6E10 species or activity-dependent presynaptic/synaptic plasticity changes^8^. This spatial pattern mirrors electrophysiological findings in 5xFAD mice describing early presynaptic dysfunction followed by postsynaptic receptor dysregulation and synaptic weakening^10^.

Isolating specific clusters with a 90% similarity threshold signifies that their structural, spatial, and molecular properties closely resemble one another. Colocalization patterns at this specificity level can reveal functional synaptic contact sites with minimal noise from divergent or atypical cluster morphologies. Aβ6E10 clusters with 90% similarity may identify amyloid populations into oligomeric versus plaque-type aggregates. Analyzing colocalization with RIM1 or GluA2 at this granularity allows identification of which amyloid species preferentially associate with presynaptic zones or postsynaptic receptors (Fig.3). These results reveal that select conformational states of RIM1 and GluA2 correspond to those seen in clusters with Aβ6E10, suggesting that Aβ6E10 selectively interacts with receptors based on their conformational states. Furthermore, 71% of Aβ6E10–RIM1 colocalizations are also triple colocalizations with GluA2, while only 1% of Aβ6E10–GluA2 colocalizations are also triple colocalizations (Table 4), meaning majority of amyloid bound to RIM1 is also bound to GluA2, forming the disease emergent ternary assembly. This asymmetry largely reflects amyloid’s near-universal GluA2 contact (∼90% of amyloid clusters), evidenced by its association with multiple GluA receptor subtypes. Further beyond that baseline, the ternary assembly is significantly enriched over independent pairwise rates (1.12×, permutation p = 0.013; Table 5), indicating that it is associated with the amyloid bridge. These findings highlight that the formation of the ternary cluster is likely enriched presynaptically, with Aβ6E10 acting as a bridge to postsynaptic GluA2, introducing a possible mechanism of synaptic disruption by Aβ6E10 trans-synaptically. Overall, our findings suggest that the observed interaction between Aβ6E10, RIM1, and GluA2 represents a mechanism of Aβ6E10 mediated synaptic dysfunction in AD, driven by Aβ6E10 interaction with specific pre- and post-synaptic conformational states.

To further explore the mechanism of synaptic dysfunction driven by our novel discovery, we focused on the nanodomain clusters and triple colocalization results. Two nanodomain clusters represent spatial proximities between two distinct protein nanodomains, often signifying direct or near-direct molecular interactions or functional coupling at synapses. These interactions modulate synaptic efficacy, plasticity, and signal transduction, which are critical to normal CNS function. However, double colocalizations represent pairwise interactions that may occur broadly across different brain regions, which could reduce therapeutic precision. Triple colocalizations pinpoint molecular microdomains where multiple pathways converge, increasing specificity for intervention compared to individual proteins. These may signify more complex molecular architectures within synapses that coordinate multiple signaling pathways simultaneously.

The presence of triple nanocluster intersections likely marks sites of enhanced signaling integration, potentially corresponding to areas of heightened synaptic plasticity or pathology in the 5xFAD model of AD. These ternary assemblies offer more precise, biologically relevant druggable interfaces, as they capture the molecular context essential for targeting multi-protein complexes contributing to pathology, rather than single proteins. When triple colocalizations include extracellular matrix (ECM) marker Lectin and glial markers (SMI for neuronal elements, GFAP for astrocytes), these likely signify specialized tripartite synapses or microenvironments where neurons, extracellular matrix, and astrocytes interact closely. In AD pathology, such assemblies may reflect reactive gliosis and ECM remodeling associated with synaptic dysfunction^16^.

The Aβ6E10-GluA2–RIM1 ternary assembly highlights key intersections between Amyloid-beta pathology and synaptic function. Aβ plaques in AD disrupt synaptic organization by altering the spatial distribution of critical proteins. This disruption at pre- and postsynaptic sites contributes to synaptic dysfunction and cognitive decline. Amyloid-beta’s synaptotoxic effects are marked by interactions with synaptic proteins and are crucial in AD pathogenesis, indicated by interference in synaptic signaling, highlighting amyloid-beta’s role in neurodegeneration^21,22^.

Triple colocalizations represent precise nanodomain assemblies, rather than widespread occurrences. The identification of the cluster highlights a mechanistic interface involved in the pathology of disease emergent synaptic dysfunction. Importantly, therapeutics designed to bind at these triple interfaces may selectively more effectively restore synaptic function or prevent degeneration.

To further validate ASCEND’s findings, we conducted a literature review to explore the plausibility of the observed GluA2-RIM1-Aβ6E10 spatial interaction. This process enabled the proposal of a likely mechanism by which these proteins interact, supporting the validity of ASCEND’s novel finding. GluA2 is composed of R1 and R2 subunits that mimic a clamshell, R1 acting as a lid and R2 as a floor. ASCEND’s unique ability to observe molecular structures in dynamic and pathological states has allowed us to identify an AD-specific gap in GluA2 R2 dimer anchoring^23^. The R2 floor of GluA2 consists of two parallel domains and a hydrophobic core that work to anchor the receptor in place. Amyloid beta consists of a hydrophobic C-terminus that plays a large role in Aβ6E10 oligomerization and toxicity^24^. ASCEND analysis suggests that Aβ6E10 oligomers may wedge themselves between the two parallel domains of R2 via competitive hydrophobic displacement, forcing them apart and destabilizing the receptor. The separation of the R2 subunits initiates a cascade that forces R1 to collapse, opening the ion pore and allowing for unregulated calcium influx.

Given that 71% of Aβ6E10-RIM1 contacts represent ternary clusters compared to 1% of Aβ6E10-GluA2 contacts (Table 4), it is likely that the interaction between Aβ6E10 and RIM1 precedes and stabilizes the disruption of GluA2. RIM1 has been observed as a regulator of presynaptic activity via syntaxin activation and subsequent vesicle fusion for NT release^25^. By anchoring NT vesicles as well as preventing voltage-gated calcium channel (VGCC) inactivation, RIM1 plays an important role in facilitating NT release^26^. In addition, RIM1 acts as an anchor for transynaptic bridges, for example anchoring presynaptic neurexin which binds to postsynaptic Neuroligin-1 or LRRTM2^25–28^. In our ASCEND scan of the 5xFAD cortex, the colocalization between GluA2 and RIM1 was measured at 18.5 nm [coloc_GluA2_RIM1_1]. This distance is the molecular signature of the synaptic cleft, suggesting “leaky” receptors may be structurally maintained in the synapse via RIM1 synaptic bridging. Importantly, defects in the bridging from Neurexin to Neuroligin-1 have been linked to synaptic dysfunction and autism^27, 28^, and Amyloid-beta interaction with both neurexin and neuroligin is associated with synaptic damage and memory loss in mice^29^. Our findings propose that RIM1 mediated synaptic bridging to postsynaptic molecules such as Neuroligin-1 holds the GluA2-Aβ6E10 complex in place, preventing the leaky GluA2 from being cleared from the synapse by endocytosis and resulting in excitotoxicity. Exploring the mechanistic underpinnings of our identified pathological interface has further enabled the in-silico design of candidate compounds targeting this pathological interface for prophylaxis and rescue, which await experimental validation.

In summary, ASCEND analysis of the 5xFAD mouse has identified a novel triple co-localization relevant to amyloid beta induced excitotoxicity in AD, and further mapped the interface to identify relevant and druggable targets.

Mechanistically, our integration with AI-driven literature mining via ASCEND^TM^ consolidated these observations with emerging insights into mitochondria-related presynaptic dysfunction and neuroinflammatory cascades that modulate synaptic integrity in 5xFAD and human brains^30^. This combined approach accelerates mechanistic understanding and pinpointing precise molecular targets for therapeutic intervention.

Despite its transformative capabilities, ASCEND^TM^’s performance is contingent on the quality and completeness of multiExR microscopy data, which can be affected by tissue processing variability and antibody accessibility. Additionally, nanoscale spatial proximity inferred by the ASCEND^TM^ engine does not confirm direct molecular interactions or functional causality, underscoring the need for rigorous experimental validation. The spatial resolution limits of multiExR (∼25–40 nm) and the co-localization thresholds may overlook finer or transient interactions critical to synaptic function. This study was limited in the number of wild type ages to compare to. Moreover, reliance on existing literature for mechanistic context may bias target prioritization toward well-characterized pathways, potentially limiting discovery of novel mechanisms.

This work, however, showcases the unprecedented capability of an AI-powered autonomous science system to transcend prior analytical limits on multiplexed nanoscale proteomic data. This tool is crucial to reveal an expansive network of novel protein spatial associations critical to understanding AD synaptic pathology^31^. Integrating raw spatial measurements with cross-domain biological knowledge not only validates known disease mechanisms, but also proposes novel, biologically plausible targets for drug development that may otherwise be missed. Future studies and applications of spatial AI systems should explore integrating multiExR with in vivo imaging, electrophysiological measures, and single-cell transcriptomics to capture dynamic synaptic remodeling and validate causality. Additionally, bridging findings from mouse models to human CNS pathology remains essential for translational relevance. Collectively, this autonomous scientific paradigm promises to revolutionize CNS therapeutic discovery by accelerating identification and validation of molecular targets for drug development.

In summation, the close double co-localizations of Aβ6E10 with presynaptic RIM1 domains, alongside GluA2 postsynaptic receptor spatial organization, implicates dysfunctional synaptic nanoarchitecture in early AD pathogenesis. Notably, distinct Amyloid species appear to preferentially associate with presynaptic zones or postsynaptic receptors, suggesting differential mechanisms by which Amyloid pathology disrupts synaptic function across these compartments. For CNS drug discovery, double colocalizations provide accessible targets with manageable complexity but may lack the functional and pathological specificity that triple or multi colocalizations offer. Conversely, triple or higher-order colocalizations enable targeting biologically integrated and disease-relevant molecular hubs, promising higher therapeutic efficacy at the cost of increased design complexity and validation challenges.

We employed an autonomous spatial AI-driven system to provide a comprehensive nanoscale spatial characterization of Amyloid-beta and synaptic nanodomains in the 5xFAD mouse model of AD via multiExR microscopy to uncover thousands of novel protein associations and defined new druggable synaptic targets using these approaches. Beyond automating literature synthesis, this system executes closed-loop scientific discovery through identifying, contextualizing, and experimentally advancing CNS therapeutic candidates. Future studies should employ AI-driven approaches to explore multiExR with in-vivo imaging and human AD tissue.

## Experimental Section

### Data Acquisition

The raw multiplexed expansion revealing microscopy datasets of the 5xFAD and WT mouse somatosensory cortex from Kang, J., Schroeder, M.E., Lee, Y. et al. 2024 available at https://dataverse.harvard.edu/citation?persistentId=doi:10.7910/DVN/JJBULY with amyloid beta (Aβ, 6E10 antibody), presynaptic RIM1, postsynaptic GluA2, and reference markers. Nine 5xFAD and eight wild-type cortical regions were analyzed. Analyses used two animals per genotype (5xFAD: samples S1 and S2; wild-type: samples S3 and S4), each contributing four to five cortical regions, for a total of nine 5xFAD and eight wild-type regions.

### Segmentation and Spatial Calibration

Channels are loaded format-agnostically (OME-Zarr / ND2 / TIF) and segmented per channel by DBSCAN with voxel-aware ε (the user-facing ε is translated to a calibrated physical distance and clamped to ≥ 4× the smallest voxel) followed by an interquartile-range core-density filter that rejects diffuse, non-cluster-like points. Intensity thresholding is auto-derived (p99.9) unless overridden.

To restrict the amyloid channel to bona fide amyloid-β and exclude non-specific antibody signals, 6E10 (Aβ6E10) clusters were subjected to a concordance filter requiring agreement across independent amyloid antibodies. The 6E10 antibody produces a diffuse background carpet in addition to labeling genuine amyloid, which would otherwise inflate amyloid colocalization counts with spurious associations. To remove this background, a 6E10 cluster was retained only if it co-localized (within 100 nm pre-expansion) with at least one D54D2 cluster and at least one 12F4 cluster, two antibodies that independently label amyloid-β. This triple-positive requirement ensures that every scored amyloid cluster is corroborated by three concordant antibodies rather than a single, potentially non-specific marker. In 5xFAD tissue the filter retained dense, plaque-associated amyloid (3,299 clusters), whereas in wild-type tissue, which lacks amyloid pathology, it retained only 51 sparse background clusters with no plaque-scale clusters. Wild-type therefore serves as an amyloid-negative control, and amyloid colocalization is reported in 5xFAD only, providing a built-in specificity check on every amyloid association in this study.

To exclude DBSCAN merge artifacts (single clusters of tens to hundreds of thousands of voxels that do not correspond to biological puncta), clusters exceeding 50,000 voxels were removed prior to colocalization for all synaptic and amyloid channels; the structural marker channel (Lectin, SMI, GFAP) was exempt because it labels genuinely large vascular and glial structures.

Voxel calibration was 162.5 × 162.5 × 250 nm post-expansion, corresponding to 9.03 × 9.03 × 13.89 nm pre-expansion at the 18× expansion factor, and was propagated through every distance metric in the pipeline.

### AI-Driven Discovery

ASCENDTM processed raw 3D protein nanocluster coordinates and morphometrics to identify spatial protein associations through voxel-based spatial hashing. Colocalization was determined by grid cell adjacency: two clusters were considered colocalized when their spatial grids contained overlapping or neighboring cells. Grid cell adjacency accommodates the ∼25 to 40 nm effective spatial resolution and ∼25 nm registration error inherent to multi-round multiplexed expansion microscopy.³²,³³

A 100 nm biological distance cutoff balances capturing likely direct or near-direct molecular interactions at synaptic sites while minimizing false positives due to random chance or spatial noise. Pairs are screened by voxel-based spatial hashing, then admitted as colocalizations when (1) their cluster grids share or neighbor a cell, or (2) their centroid-to-centroid distance is ≤ 100 nm pre-expansion. Two metrics are reported per event: the grid-overlap fraction (overlapping cells divided by the smaller cluster’s cells) and the centroid distance. Distributions of distances, overlap fractions, cluster volumes, and colocalization event counts were computed using Eratos AI.

### Statistical Testing

For each channel pair in each region, statistical significance was assessed by a density-matched label-shuffling permutation test (n = 500). The null was constructed by re-permuting the two channels’ labels among that pair’s pooled clusters only, preserving the combined spatial density of the two channels, and recomputing the colocalization count under the identical detection rule applied to the observed data. This per-pair density matching prevents sparse markers from being diluted against the full cluster population, which would otherwise inflate enrichment. Each permutation used a deterministic per-pair seed, so all enrichment ratios and p-values are exactly reproducible. Per-pair enrichment was computed as the observed count divided by the mean null count, with the raw p-value, the mean and standard deviation of the null, and a 95% null confidence interval (2.5 to 97.5 percentile) reported. Benjamini-Hochberg FDR correction (alpha = 0.05) was applied across all channel pairs in the amyloid–synaptic analysis panel (28 pairs from the eight markers Aβ6E10, D54D2, 12F4, RIM1, and GluA1–GluA4; 24–28 per region depending on cluster presence), so the reported synaptic and amyloid pairs are significant after correction across the full panel rather than in isolation.

Within-condition significance is reported as the number of regions in which a pair is significantly enriched (out of 9 5xFAD and 8 wild-type). Between-genotype differences were tested by Mann-Whitney U on per-region enrichment ratios for pairs present in both genotypes.

To test whether three-way assemblies form more often than expected from the constituent pairwise rates, we applied a density-matched ternary permutation test. Within each region, GluA2-contact labels were randomly reassigned across the amyloid cluster pool, preserving the per-region GluA2-contact count, and the number of amyloid clusters contacting both a RIM1 and a GluA2 cluster was recomputed. 20,000 permutations were pooled across the nine 5xFAD regions, with a deterministic seed for reproducibility. Observed bridges were compared to this null to obtain an enrichment ratio and a one-sided p-value.

The grid-based spatial overlap criterion for defining colocalization events was chosen to capture meaningful molecular proximities within synaptic nanodomains, which typically span tens to hundreds of nanometers. This approach balances specificity (requiring actual spatial intersection of cluster volumes) with sensitivity (allowing for biological variability and minor registration artifacts inherent to multi-round multiplexed expansion microscopy).

### Statistical Validation of Colocalizations

Before any colocalization is reported, ASCEND applies a sequence of statistical and structural gates. Per-channel nanoclusters are segmented from the raw multiExR volumes by DBSCAN with a voxel-aware ε (translated to a calibrated physical distance and clamped to at least 4× the smallest voxel), an interquartile-range core-density filter, and an automatic 99.9th-percentile intensity threshold; amyloid (6E10) clusters are further restricted to those concordant with both D54D2 and 12F4, and clusters exceeding 50,000 voxels (excluding the structural Lectin, SMI, GFAP channel) are removed as merge artifacts. Voxel calibration of 162.5 × 162.5 × 250 nm post-expansion (9.03 × 9.03 × 13.89 nm pre-expansion at the 18× expansion factor) is propagated through every distance metric.

Two clusters are scored as colocalized when (1) their voxel-hashed grids share or neighbor a cell, or (2) their centroid-to-centroid distance is ≤ 100 nm pre-expansion. For each channel pair, a density-matched label-shuffling permutation test (n = 500) is then performed: the two channels’ labels are re-permuted among that pair’s pooled clusters only, preserving their combined spatial density, and the colocalization count is recomputed under the same detection rule used on the observed data, with a deterministic per-pair seed ensuring exact reproducibility. Per-pair raw p-values, the mean and standard deviation of the null, and a 95% null confidence interval (2.5 to 97.5 percentile) are reported, with Benjamini-Hochberg FDR correction (alpha = 0.05) applied across pairs and within-condition significance summarized as the number of significantly enriched regions. Between-genotype contrasts (5xFAD vs wild-type) use a Mann-Whitney U test on per-region enrichment ratios.

Cluster-to-cluster structural similarity (used to gate the ≥ 90% homogeneity selections in Figure 3) is computed as cosine similarity (≥ 0.90) in a self-supervised PointJEPA point-cloud embedding space trained on 3D ExM point clouds, a learned metric over nanocluster geometry evaluated cross-channel without channel-identity leakage rather than a heuristic shape descriptor. Validation of the resulting drug-target nominations is performed downstream of identification: each candidate cluster is mapped to a PDB structure, or for interaction-based designs to a Chai-1-predicted complex, and pocket druggability, volume, and residue composition are scored by fpocket, with the top-druggability pocket selected unless explicitly overridden. Designed peptides, nanobodies, and antibodies are generated with BoltzGen on H100 GPUs; small-molecule designs are produced by DiffSBDD, followed by AlphaFold-multimer modeling, Boltz-ternary scoring, OpenMM energy minimization, and cryptic-pocket discovery. All candidates are then filtered for pan-assay interference (PAINS) liability, heavy-atom limits, and structural novelty against the PDB via Foldseek; any design that fails a filter is surfaced as a rejected candidate with a reason code rather than silently dropped.

Together, these layers, statistical (permutation testing with FDR correction), structural (self-supervised embedding similarity), and design-side (Chai-1, fpocket, BoltzGen, DiffSBDD, Foldseek, PAINS), constitute the in-silico validation that gates which targets and which designs leave the platform. Wet-lab biochemical confirmation of direct binding (Co-IP, SPR) and functional rescue experiments are scoped as the next, prospective phase of the closed-loop pipeline and are not claimed in the present study.

### Integration with Biomedical Literature

ASCEND^TM^ cross-referenced and discovered spatial associations against a comprehensive biomedical knowledge graph constructed from PubMed, Nature portfolios, and neurobiological databases. The ASCEND^TM^ engine has access to all of Pubmed’s 39 million articles and clinicaltrials.gov, which comprises 500,000 trials to date. Through leveraging natural language processing and mechanistic ontologies, Eratos AI identified associations from thousands of scientific articles that demonstrated significant concordance with validated disease mechanisms, while flagging novel associations for further exploration (Supplemental Table 1).

### Mechanistic Synthesis and Target Prioritization

Spatial patterns and literature-derived mechanisms were integrated into mechanistic network models elucidating candidate drug targets’ roles in synaptic pathology and disease progression. ASCEND^TM^ evaluated binding interface accessibility, target specificity, and pathway centrality to prioritize targets with high translational potential. The analyzed 5xFAD and wild-type somatosensory cortex datasets included multiple imaged protein targets and markers, comprising the AMPA-receptor subunits GLUA1, GluA2, and GLUA3, the presynaptic active-zone protein RIM1, amyloid-β (Aβ6E10), and reference markers (Lectin, SMI, GFAP). Double colocalizations represent spatial proximities between two distinct protein nanodomains, often signifying direct or near-direct molecular interactions or functional coupling at synapses. Triple colocalizations involve spatial overlap or close proximity of three distinct protein nanodomains or markers, representing highly integrated molecular assemblies

### Accelerating Scientific Discovery

To quantify the acceleration achieved by ASCEND^TM^ relative to conventional human-driven analysis, we benchmarked the system’s performance on key closed-loop workflows including nanoscale cluster segmentation, pairwise colocalization analysis, multi-dimensional morphometric comparison, literature integration, and hypothesis generation. Across 17 multiExR volumetric regions (9 5xFAD and 8 wild-type), ASCEND^TM^ autonomously segmented 161,678 clusters, computed over 665 million pairwise spatial relationships, and cross-referenced 4,391 scientific papers using parallelized algorithms and high-throughput natural language processing. Overall, ASCEND^TM^ was able to locate thousands of colocalizations in a cubic micrometer of raw multiplexed scans in minutes, understand the spatial context from the raw scans to filter down millions of journal articles and papers, and connect its ground truth to that filtered literature in seconds. By comparison, equivalent manual analysis by expert scientists would require weeks to months given the complexity and volume of spatial data and breadth of relevant literature. The AI’s iterative capability to rapidly cycle between data extraction, contextualization, hypothesis refinement, and experimental design enables true closed-loop discovery, accelerating insight generation.

## Associated Content

### Supporting Information

**Table S1 | Representative Publications Used for Literature Integration and Novel Findings Contextualization**

## Ancillary Information

**Authors:**

Maya Viviana Suissa

3915 Dixie Canyon Avenue

Sherman Oaks, California 91423

213-407-0081

Anne Koutures

1226 6th Ave.

San Francisco, CA 94122

310-744-5686

Samantha Harker

427 E Tyler Mall

Tempe, AZ 85281

213-222-7948

Brent Vaughan

445 Ramona Road

Portola Valley, CA 94028

650-207-3741

Jonathan Kfir

7 Nidden

Irvine, CA 92603

949-943-4367

Anoushka Bhat

410 West 53rd Street, Apt 419

New York, New York 10019

909-345-4385

Ryan Shihabi

1110 Town and Country Rd, Unit 561

Orange, CA 92868

949-505-4300

Mahender Dewal

77 Massachusetts Ave.

Cambridge, MA. 02139

803-397-0383

Edward Boyden

77 Massachusetts Ave.

Cambridge, MA., 02139

617-324-3085

## Acknowledgements

We thank Dr. Allan Levey and the anonymous reviewers for valuable feedback in reviewing this manuscript.

## Abbreviations

5xFAD: Five familial Alzheimer’s disease mutations; Aβ6E10: Amyloid-beta 6E10 antibody; AD: Alzheimer’s disease; AMPAR: α-amino-3-hydroxy-5-methyl-4-isoxazolepropionic acid receptor; ASCEND: AI for Spatial Computing and Embedded Neurotherapeutic Discovery; ECM: Extracellular matrix; ExM: Expansion microscopy; GFAP: Glial fibrillary acidic protein; GluA2: Glutamate ionotropic receptor AMPA type subunit 2; MAPT: Microtubule-associated protein tau; multiExR: Multiplexed expansion revealing; NDC: N-Dimensional Colocalization; RIM1: Rab3-interacting molecule 1; ROI: Region of interest; S1ROI1: Somatosensory cortex region of interest 1 (5xFAD); S3ROI1: Somatosensory cortex region of interest 1 (Wild Type); SMI: Neurofilament marker; synGAP: Synaptic GTPase-activating protein.

