## Supplemental Table 1 for "Autonomous AI-Driven Nanoscale Spatial Mapping Reveals Novel Targets and Ternary Architectures in 5xFAD Alzheimer’s Disease Model"

1) Eratos Therapeutics
2) University of California - San Francisco, School of Pharmacy
3) Arizona State University, School of Life Sciences
4) Chapman University, School of Engineering
5) Massachusetts Institute of Technology, Department of Chemistry
6) Massachusetts Institute of Technology, Department of Biological Engineering
7) Massachusetts Institute of Technology, McGovern Institute
8) Massachusetts Institute of Technology, Koch Institute
9) Massachusetts Institute of Technology, Center for Neurobiological Engineering
10) Massachusetts Institute of Technology, K. Lisa Yang Center for Bionics and Yang Tan Collective
11) Massachusetts Institute of Technology, Department of Brain and Cognitive Sciences
12) Rady Children's Health Orange County, Sharon Disney Lund Medical Intelligence, Information, Investigation, and Innovation Institute
13) University of California - Irvine, School of Medicine, Department of Pediatrics
14) University of California - Riverside, School of Medicine, Department of Psychiatry & Neurosciences

**Table S1 | Representative Publications Used for Literature Integration and Novel Findings Contextualization**

| Topic | Representative Key Publications | Relevance to Study |
|---|---|---|
| Alzheimer's disease synaptic dysfunction | Selkoe DJ. Alzheimer's disease is a synaptic failure. *Science*. 2002;298(5594):789-791. https://doi.org/10.1126/science.1074069 | Foundational review on synaptic pathology in AD |
| Amyloid beta oligomers and synapse | Li S, Hong S, Shepardson NE, et al. Soluble oligomers of amyloid beta protein facilitate hippocampal long-term depression by disrupting neuronal glutamate uptake. *Neuron.* 2009;62(6):788-801. | Mechanistic study on Aβ oligomers disrupting synaptic activity |
| 5xFAD mouse model pathology | Oakley H, et al. Intraneuronal beta-amyloid aggregates, neurodegeneration, and neuron loss in transgenic mice with five familial Alzheimer's disease mutations. *J Neurosci.* 2006;26(40):10129-40. | Key model characterization underpinning our spatial data source |
| Presynaptic active zone and RIM1 | Schoch S, et al. RIM1alpha forms a protein scaffold for regulating neurotransmitter release at the active zone. *Nature.*2002;415(6869):321-6. | Molecular mechanisms of RIM1 in neurotransmission |

| | | |
|---|---|---|
| Expansion microscopy and nanoscale imaging | Chen F, Tillberg PW, Boyden ES. Expansion microscopy. *Science.* 2015;347(6221):543-548. | Basis for multiExR imaging technology |
| Multiplexed expansion revealing technique | Kang J, Schroeder ME, Lee Y, et al. Multiplexed expansion revealing for imaging multiprotein nanostructures in healthy and diseased brain. *Nat Commun.*2024;15:9722. | Methodological foundation and source of raw spatial proteomic data |
| SynGAP and Alzheimer's disease synapses | G. Rumbaugh, J.P. Adams, J.H. Kim, & R.L. Huganir, SynGAP regulates synaptic strength and mitogen-activated protein kinases in cultured neurons, Proc. Natl. Acad. Sci. U.S.A. 103 (12) 4344-4351, https://doi.org/10.1073/pnas.0600084103 (2006). | Illuminates role of synGAP, connected to synaptic remodeling discovered in present study |
| Neuroinflammation in AD and synaptic loss | Heneka MT, et al. Neuroinflammation in Alzheimer's disease. *Lancet Neurol.*2015;14(4):388-405. | Supports inflammatory pathways related to molecular targets identified |
| Mitochondrial dysfunction in AD synapses | Wang X, Su B, Lee HG, Li X, Perry G, Smith MA, Zhu X. Impaired balance of mitochondrial fission and fusion in Alzheimer's disease. J Neurosci. 2009 Jul 15;29(28):9090-103. doi: 10.1523/JNEUROSCI.1357-09.2009. PMID: 19605646; PMCID: PMC2735241. | Adds context for mechanistic pathways integrated via AI |

| | | |
|---|---|---|
| Synaptic plasticity impairments in AD | Shankar GM, et al. Amyloid-beta protein dimers isolated directly from Alzheimer's brains impair synaptic plasticity and memory. *Nat Med.* 2008;14(8):837-842. | Functional evidence supporting the synaptic disruption linked to spatial protein data |
